# Molecular cloning and host range analysis of three cytomegaloviruses from *Mastomys natalensis*

**DOI:** 10.1101/2024.12.05.626976

**Authors:** Laura Staliunaite, Olha Puhach, Eleonore Ostermann, Kyle Rosenke, Jenna Nichols, Lisa Oestereich, Heinz Feldmann, Andrew J. Davison, Michael A. Jarvis, Wolfram Brune

## Abstract

Herpesvirus-based vectors are attractive for use as conventional or transmissible vaccines against emerging zoonoses in inaccessible animal populations. In both cases, cytomegaloviruses as members of the subfamily *Betaherpesvirinae* are particularly suitable for vaccine development as they are highly specific for their natural host species, infect a large proportion of their host population, and cause mild infections in healthy individuals. The Natal multimammate mouse (*Mastomys natalensis*) is the natural reservoir of Lassa virus, which causes deadly hemorrhagic fever in humans. *M. natalensis* was recently reported to harbor at least three different cytomegaloviruses (MnatCMV1, MnatCMV2 and MnatCMV3). Herein, we report the molecular cloning of three complete MnatCMV genomes in a yeast and bacterial artificial chromosome (YAC-BAC) hybrid vector. Purified viral genomes were cloned in yeast by single-step transformation-associated recombination (STAR cloning) and subsequently transferred to *Escherichia coli* for further genetic manipulation. Integrity of the complete cloned viral genomes was verified by sequencing, and replication fitness of viruses reconstituted from these clones was analyzed by replication kinetics in *M. natalensis* fibroblasts and kidney epithelial cells. We also found that neither parental nor cloned MnatCMVs replicated in mouse and rat fibroblasts, nor did they show sustained replication in baby hamster kidney cells, consistent with the expected narrow host range for these viruses. We further demonstrated that an exogenous sequence can be inserted by BAC-based mutagenesis between open reading frames M25 and m25.1 of MnatCMV2 without affecting replication fitness *in vitro*, identifying this site as potentially suitable for the insertion of vaccine target antigen genes.

**Importance:** Cytomegaloviruses recently discovered in the Natal multimammate mouse (*Mastomys natalensis*) are widespread within the *M. natalensis* population. Since these rodents also serve as natural hosts of the human pathogen Lassa virus (LASV), we investigated the potential suitability of *M. natalensis* CMVs (MnatCMVs) as vaccine vectors. We describe the cloning of three different MnatCMV genomes as bacterial artificial chromosomes (BACs). Replicative capacity and species specificity of these BAC-derived MnatCMVs were analyzed in multiple cell types. We also identified a transgene insertion site within one of the MnatCMV genomes suitable for the incorporation of vaccine target antigens. Together, this study provides a foundation for the development of MnatCMVs as transmissible MnatCMV-based LASV vaccines to reduce LASV prevalence in hard-to-reach *M. natalensis* populations and thereby zoonotic transmission to humans.

## Introduction

*Orthoherpesviridae* is a diverse family of large double-stranded enveloped DNA viruses that establish lifelong infection within their host. These viruses are generally highly host specific with different family members infecting a wide range of different host species, including humans (1). Human cytomegalovirus (HCMV) belongs to the *Betaherpesvirinae* subfamily and is highly prevalent in the human population. HCMV causes mild infections in the immunocompetent host, but can cause severe disease in immunocompromised individuals, such as transplant recipients and newborns (2). Although HCMV is restricted to humans (3), many animals harbor their own specific cytomegaloviruses (CMVs), such as mouse CMV (MCMV), rat CMV (RCMV), guinea pig (GPCMV), and rhesus macaque CMV (RhCMV) (4). Recently, four novel CMVs were identified in wild Natal multimammate mice (*Mastomys natalensis; M. natalensis*) from Mali and Cote d’Ivoire (5). Full genome sequencing of three different *M. natalensis* cytomegaloviruses (MnatCMV1, MnatCMV2 and MnatCMV3) confirmed the close relationship of these viruses to other rodent CMVs such as MCMV and RCMV (6).

*M. natalensis* can be found in rural areas of Sub-Saharan Africa and are known to be the main host for the zoonotic arenavirus, Lassa virus (LASV). LASV continues to pose a global zoonotic threat to humans, leading to fatal hemorrhagic fever infections in endemic areas of West Africa. In addition, isolated imported cases in Europe and the USA have been documented several times in the past (7). There are currently no Lassa fever vaccines commercially available for human use. Human LASV infections are primarily attributable to spill-over events from the *M. natalensis* reservoir, as human-to-human transmissions occur only in rare cases (8). As contact between the natural host and humans is predicted to expand constantly in the future due to climate change, increased land use, and human population growth, a protection strategy against the transmission of LASV from this key species to humans is urgently required (9).

The recent discovery and genetic characterization of multiple MnatCMVs in *M. natalensis* have opened the possibility of using MnatCMVs as transmissible vaccine vectors against LASV. Using this strategy, a replication-competent MnatCMV vaccine vector carrying a LASV antigen gene would be used to transmit LASV immunity between individual *M. natalensis* animals and thereby reduce the risk of zoonotic transmission to humans. Previous studies have described and discussed the suitability of using CMVs as vaccine vectors, which is based on multiple characteristics including their ability to cause latent infections and induce strong T cell responses with long-lasting T cell memory (10–14) Substantial coding capacity of CMV potentially enables the insertion of large regions of heterologous genetic material such as vaccine antigen genes. CMVs are also known to cause only mild infection in their natural immunocompetent host, and are highly restricted to their individual host species, preventing off-species spread. Importantly for a ubiquitous virus, CMVs are also capable of super-infecting previously infected animals, suggesting that CMV-based vectors may be able to transmit irrespective of prior CMV infection history (15, 16). For instance, *M. natalensis* animals quite frequently carry multiple different MnatCMVs simultaneously (6).

Cloning of herpesvirus genomes as bacterial artificial chromosomes (BACs) in *E. coli* allows the conservation of defined molecular clones and their precise modification by BAC recombineering (17–19). The standard method for BAC cloning of herpesvirus genomes, first described by Messerle and colleagues for the cloning of an MCMV genome (20), involves the insertion of the BAC replicon into the viral genome by homologous recombination in virus-infected cells. Subsequently, circular viral genomes containing the BAC replicon are isolated and introduced into *E. coli*. In recent years, faster and more efficient BAC cloning methods have been developed that rely on transformation-associated recombination (TAR) in yeast. These TAR-based approaches allow the assembly of complete herpesvirus genomes from cloned overlapping genome fragments (21, 22) or, as we have shown recently, the cloning of entire herpesvirus genomes in a single step (STAR cloning)(23).

The present study provides the foundation for future development of MnatCMVs as vaccine vectors. Our aim was to clone complete genomes of all three MnatCMVs as BACs by STAR cloning. The integrity and completeness of these BAC-cloned MnatCMV genomes was then verified, followed by analysis of their replication fitness and host range in cell culture. Finally, an insertion site suitable for incorporating foreign genetic material was identified by genetic manipulation and characterization of one of the BAC-cloned viruses.

## Results

### Generation of an MnatCMV2 BAC by STAR cloning

Our aim was to generate BAC clones of MnatCMV type 1, 2 and 3 genomes by STAR cloning. This was first performed using the MnatCMV2 genome. This genome is comprised of a 206,952 bp long unique region flanked by 30 bp direct repeats at each terminus. Since the 30 bp terminal repeat might compromise the success of STAR cloning, we decided to clone only the long unique region. To do this, linear viral DNA (vDNA) was isolated from MnatCMV2 virions released into the supernatant of infected MasEFs. The cloning vector (pCC1BAC-his3), consisting of a yeast centromeric plasmid (YCp) replicon, a BAC replicon, and selectable markers, was modified to contain two 60 bp homology hooks identical to sequences near the end of the MnatCMV2 long unique region. The cloning vector was linearized at a ClaI restriction site located between the homology hooks (Fig. 1A). Transformation of yeast (*S. cerevisiae*) with vDNA and the linearized vector resulted in several yeast colonies growing on *his3* deficient agar plates, indicating a successful incorporation of the pCC1BAC-his3 vector carrying the *his3* selection marker. Colonies were screened by PCR for the presence of the MnatCMV2 genome using primers specific for the MnatCMV2 genome. DNA isolated from two PCR-positive yeast clones (clone 1 and 2) were then used to transform of *E. coli*. BAC DNA purified from three *E. coli* clones was analyzed by restriction digestion (Fig. 1B) and Illumina sequencing. Clones 2A and 2B (originating from the same yeast clone 2) were found to be complete and identical to the corresponding parental viral genome sequence.

**Figure 1.**
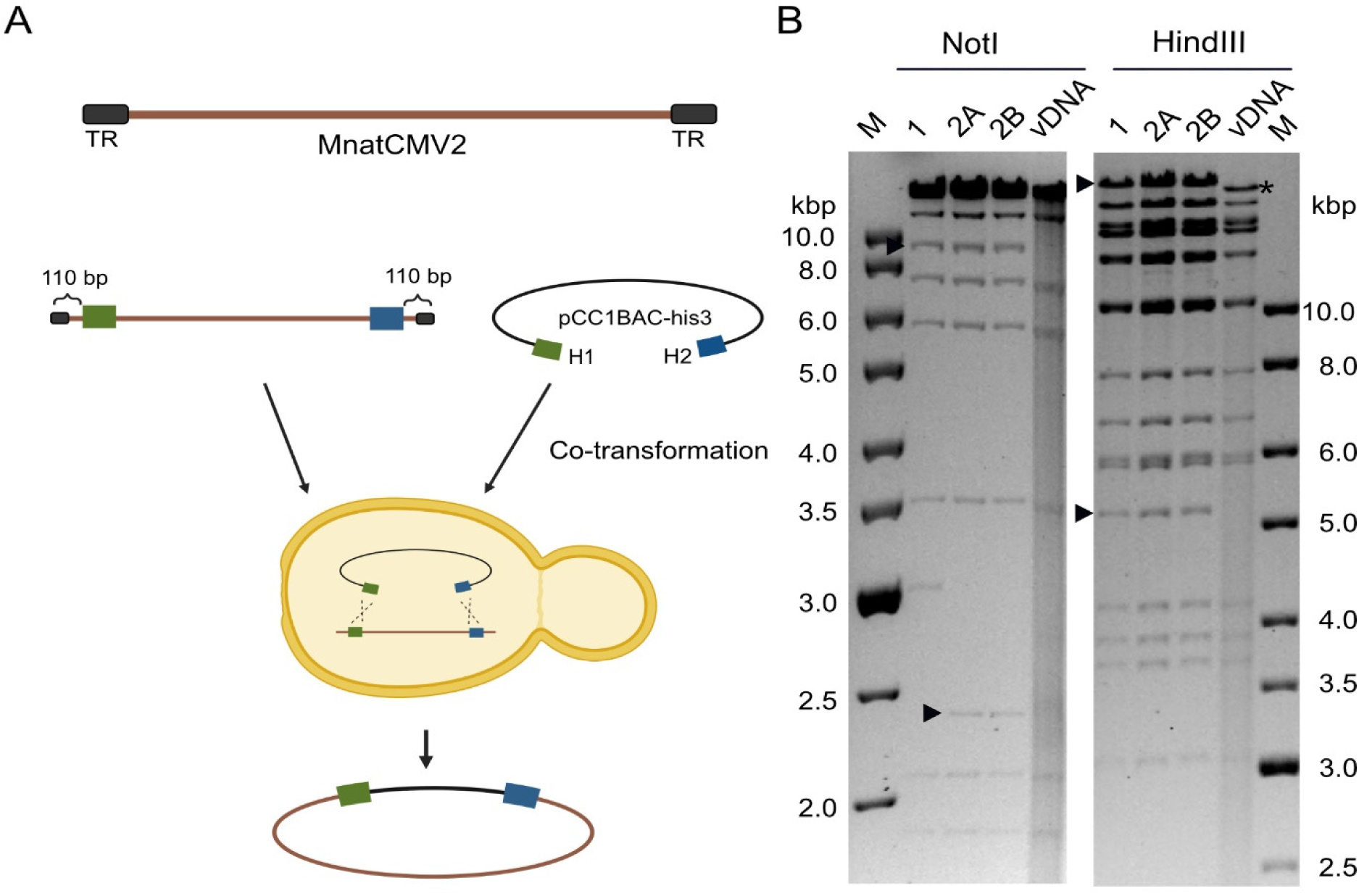
STAR cloning of MnatCMV2. **(A)** Linearized cloning vector and vDNA from MnatCMV2 virions were used to transform yeast spheroplasts. Homology hooks (60 bp, blue and green) for recombination were homologous to regions 110 bp upstream and downstream from the vDNA termini. Circular DNA consisting of the vector and the long unique region from vDNA was isolated from yeast and transferred to *E. coli*. **(B)** BAC DNA originating from two independent yeast clones isolated from *E. coli* clones was digested with NotI or HindIII and analyzed by gel electrophoresis with vDNA as a reference. M, DNA size markers. Vector-vDNA junction fragments (arrow heads) and genome terminal fragments (asterisks) absent from the circular BAC clones are indicated.

As the MnatCMV2 clones did not include the genome termini, we restored the full length genome in BAC clone 2B by re-inserting the missing sequences using *en passant* BAC mutagenesis. PCR was used to amplify the joined ends of the viral genome using DNA from MnatCMV2-infected cells as a template. The 1 kbp PCR product, containing 500 bp each from the right and left ends of the viral genome, was then cloned in pcDNA3 (pcDNA_MnatCMV_HR). This plasmid was further modified by inserting a kanamycin resistance (Kan) marker and an I-SceI restriction site (I-Sce/Kan), flanked by 80 bp of duplicated sequence for excision of the Kan marker. The resulting plasmid (Fig. 2A) was used as template for subsequent *en passant* mutagenesis steps.

**Figure 2.**
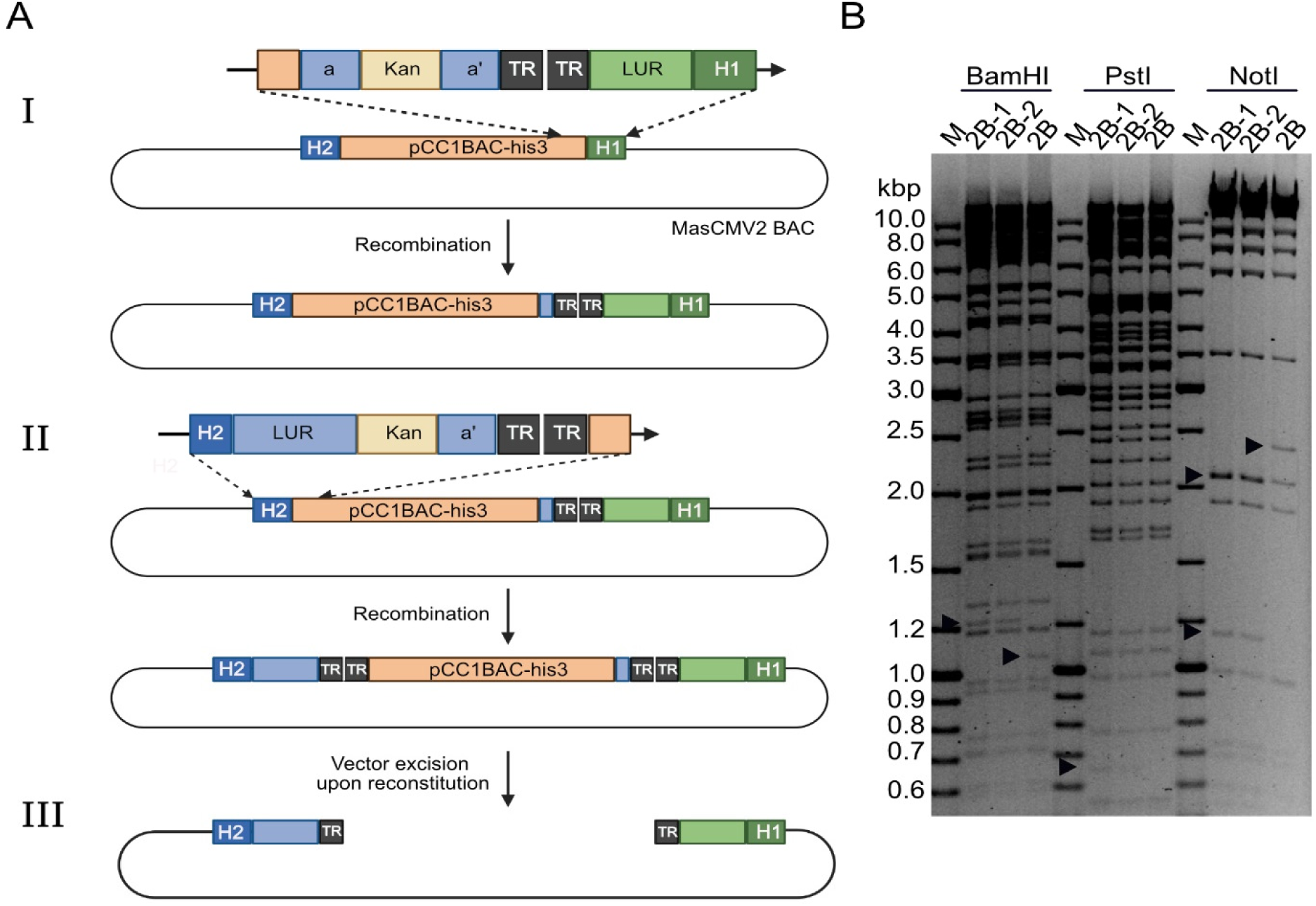
Repair of MnatCMV2 BAC via *en passant* mutagenesis. **(A)** The incomplete MnatCMV2 BAC (clone 2B), which contained the long unique region (LUR) without the terminal repeats (TR), was repaired by two rounds of *en passant* mutagenesis. (I) Repair of the left terminus of the MnatCMV2 genome using a linear DNA fragment containing the missing 80 bp of the LUR region (green), two TRs (black) and an I-Sce/Kan cassette (yellow) flanked by an 80-bp duplication (a and a’). Sequences homologous to H1 (dark green) and the pCC1BAC-his3 vector (orange) were used for recombination. The Kan cassette was subsequently removed in the second step of *en passant* mutagenesis. (II) Repair of the right terminus of the MnatCMV2 genome was performed in the same way as above. The final construct contains the full-length MnatCMV2 genome with a copy of TR at each end. (III) Complete linear MnatCMV2 genome after cleavage by the viral terminase. **(B)** Restriction fragment length analysis of BAC DNA before and after repair of the MasCMV2 genome termini. The unrepaired MnatCMV2 BAC (clone 2B) lacking the genomic termini and two repaired BAC clones (2B-1 and 2B-2) obtained after two rounds of BAC mutagenesis are shown. Arrow heads indicate genome differences resulting from insertion of left and right genome termini. M, DNA size marker.

The left junction (LJ) was repaired first. A PCR product containing the missing left end of the genome, the two terminal repeats, and the I-SceI/Kan marker was inserted by homologous recombination into the LJ of the MnatCMV2 BAC using appropriate primers (Table 1). The Kan marker was subsequently removed by a second homologous recombination step. The right junction (RJ) was repaired in an analogous fashion (Fig. 2A). The resulting BAC contained the full-length MnatCMV2 genome and two tandem TRs at either end of the genome. As the viral terminase enzyme usually cuts between the TRs, excision of the YcP-BAC cloning vector was expected to occur automatically during genome replication and packaging.

**Table 1.**
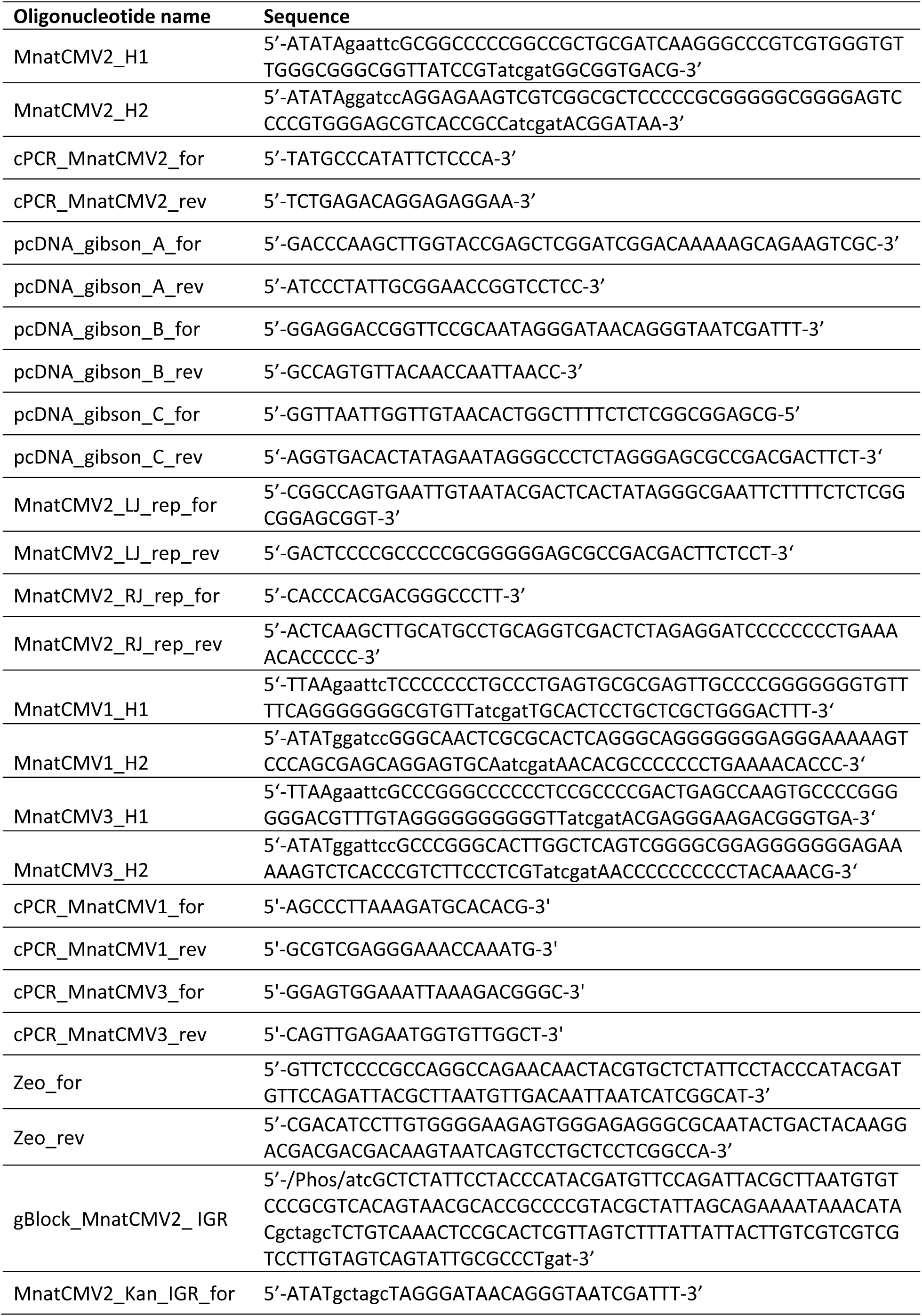
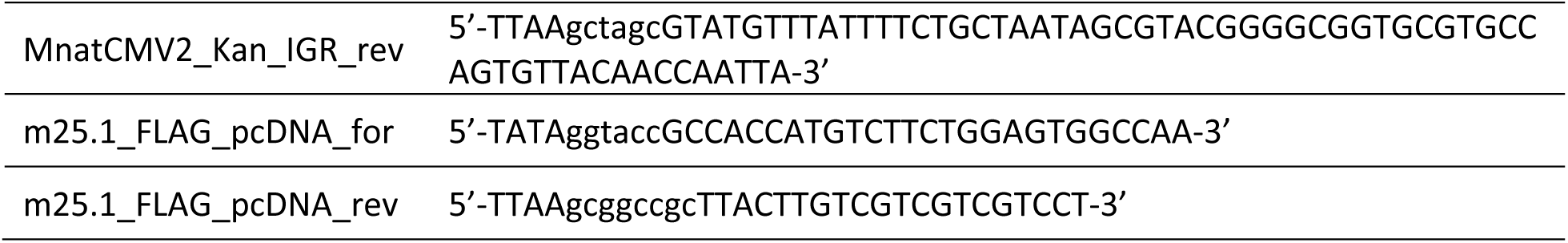
List of oligonucleotides and gBlocks used in this study. Restriction sites are shown in lower case.

Two independent clones of the repaired MnatCMV2 BAC were analyzed by restriction digest with different endonucleases (Fig. 2B). The restriction pattern matched expectations. Upon sequencing of both clones, no unintended mutations or sequence alterations were detected. To determine whether the BACs could give rise to infectious virus, MasEFs were transfected with BAC DNA. After 6 days, plaques were observed, and the infection subsequently spread through the fibroblast monolayer. To verify the integrity of the reconstituted viral genome (rMnatCMV2) and correct excision of the vector, viral DNA was isolated and subjected to Illumina sequencing. Again, no sequence alterations were detected. The genome ends each contained a single TR and were devoid of vector sequences, indicating that the YCp-BAC vector was excised as expected.

### Generation of MnatCMV1 and MnatCMV3 BACs by STAR cloning

Having successfully cloned the MnatCMV2 genome, we wanted to show the reproducibility of this approach for MnatCMVs by cloning the genomes of MnatCMV1 and MnatCMV3 as BACs. Whilst cloning MnatCMV2, we had also STAR-cloned the entire genomes (including the TRs) of two RCMV strains, but, notably, this was achieved in a single step, (i.e. without the need to re-insert the TRs by BAC recombineering (23)). For this purpose, the homology hooks contained the very first and last 60 bps of the respective viral genome, consisting of 30 bp of TR and 30 bp from the adjacent long unique region (Fig. 3A). MnatCMV1 and MnatCMV3 DNAs were isolated from viral particles and used for STAR cloning essentially as described above using this modified strategy. PCR screening of yeast colonies and transfer of cloned genomes from *S. cerevisiae* to *E. coli* were performed as described above. One complete and sequence-verified BAC clone of both MnatCMV1 and MnatCMV3 (Fig. 3B) were selected for subsequent experiments.

**Figure 3.**
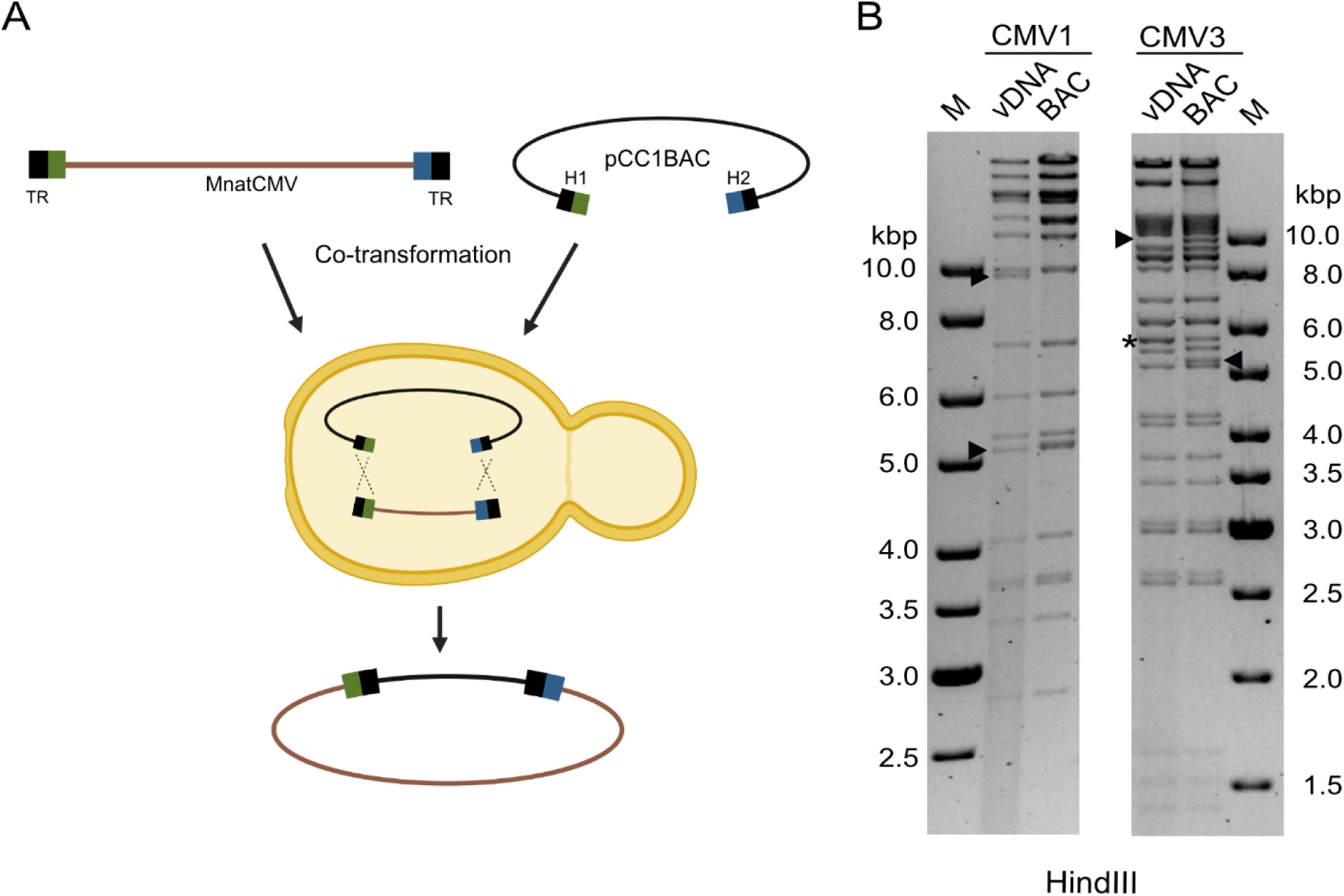
STAR cloning of MnatCMV1 and MnatCMV3. **(A)** Linearized pCC1BAC-his3 cloning vectors with homology hooks for vDNA isolated from MnatCMV1 and MnatCMV3 virions were used for transformation of yeast spheroplasts. The homology hooks consisted of the first and last 60 bp of the viral genome: 30 bp of the long unique region (green or blue) and a 30-bp terminal repeat (TR). **(B)** Restriction fragment length analysis of MnatCMV1 and MnatCMV3 BACs. vDNA is included for comparison. Vector-vDNA junction fragments (arrow heads) and terminal genomic fragments absent in the circular BAC clones (asterisks) are indicated. M, DNA size marker.

Infectious virus was reconstituted from MnatCMV1 and MnatCMV3 BACs by transfection of MasEFs (rMnatCMV1 and rMnatCMV3). Plaques were observed 6 to 8 days after transfection, and virus stocks were prepared. The integrity of the viral genome and correct excision of the vector was verified in each case by Illumina sequencing of viral DNA.

### Replication fitness and host range of reconstituted MnatCMV

To confirm further that the BAC-derived MnatCMVs are equivalent to their parental viruses, their replication was compared in MasEFs and Mastomys kidney epithelial cells (MasKECs). In multistep replication kinetics, the BAC-derived MnatCMVs replicated to titers comparable to the parental viruses (Fig. 4A and 4B).

**Figure 4.**
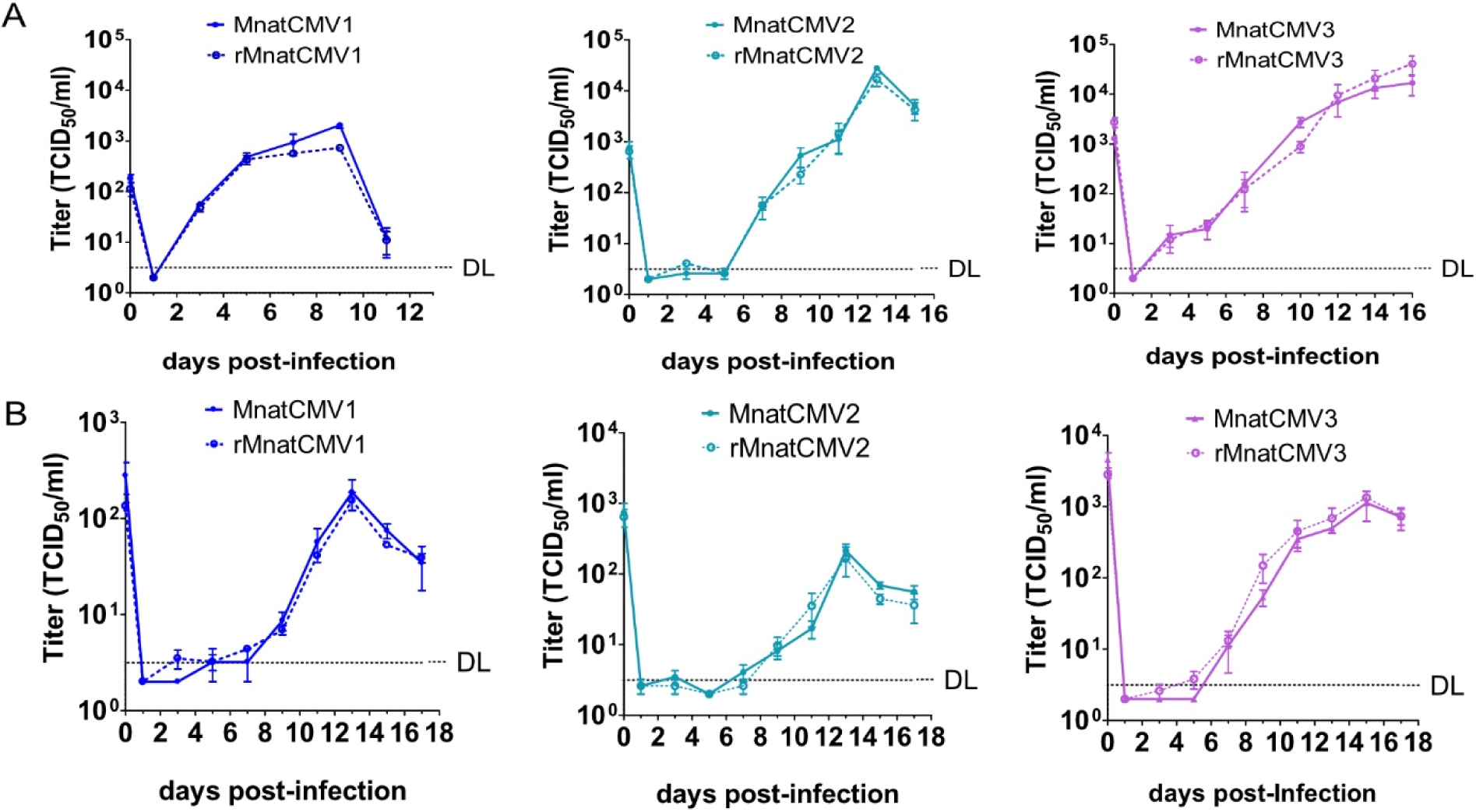
Multistep replication kinetics of MnatCMVs. **(A)** MasEFs were infected at a multiplicity of infection (MOI) of 0.1 with the parental MnatCMVs or the BAC-derived rMnatCMVs. Viral titers in the supernatants were determined. **(B)** MasKECs were infected (MOI 0.5) with the parental MnatCMVs or the BAC-derived rMnatCMVs. Viral titers in the supernatants were determined. Mean ±SEM of three biological replicates are shown. DL, detection limit.

Host species specificity is an important factor when choosing a replication-competent vaccine vector. Although CMVs are generally considered to be highly species specific, there are examples of CMVs replicating in cells from a closely related species. For instance, MCMV was reported to replicate in rat cells (24–26). Therefore, we tested whether the three MnatCMVs can replicate in fibroblasts from other rodent species. Murine NIH-3T3 fibroblasts and rat embryonic fibroblasts (REFs) were used as they are highly permissive for MCMV and RCMV, respectively. Baby hamster kidney (BHK-21) fibroblasts were also tested as these cells are permissive for many viruses. These cells were used for multistep replication kinetics with all three MnatCMVs (Fig. 5). In NIH-3T3 and REF cells, no replication was detected for all three MnatCMVs (Fig. 5A and 5B). Very low quantities of MnatCMV2 were released by BHK-21 cells, but virus release was transient and only just above the detection limit (Fig. 5C). Overall, the data indicate that the MnatCMVs are highly host cell species-specific.

**Figure 5.**
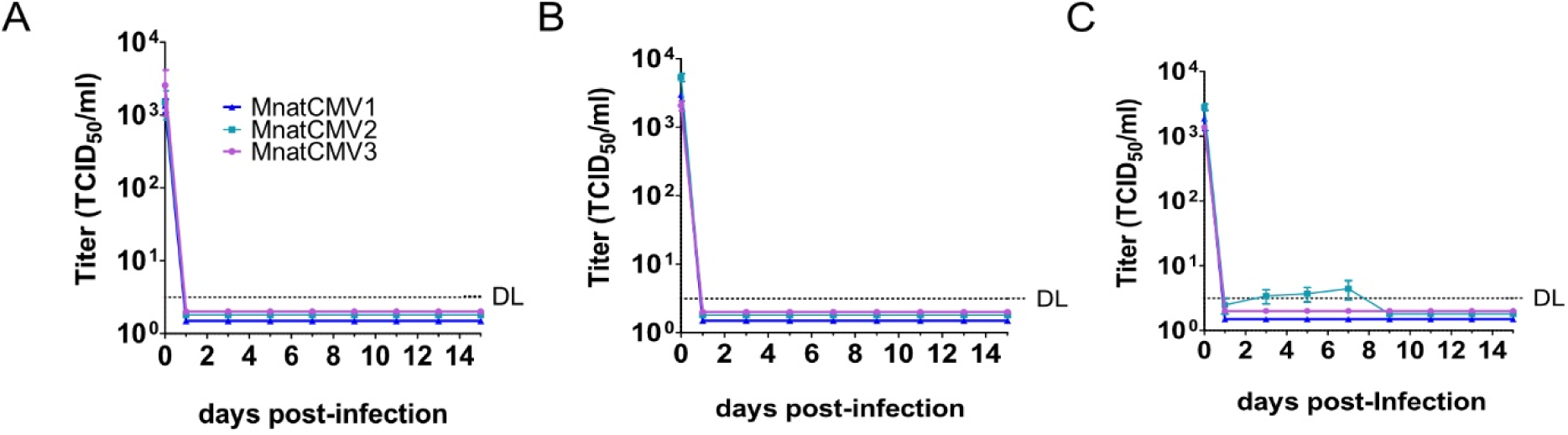
Replication kinetics of MnatCMVs in **(A)** mouse embryonic fibroblasts (NIH-3T3), **(B)** rat embryonic fibroblasts (REFs), and **(C)** baby hamster kidney fibroblasts (BHK-21). Cells were infected at an MOI of 0.5. Viral titers in the supernatants were determined. Mean ±SEM of three biological replicates are shown. DL, detection limit.

### Identification of a suitable transgene insertion site within the MnatCMV2 genome

Our final aim was to find a suitable transgene insertion site within the MnatCMV genome. For use as a transmissible vaccine, the insertion of a transgene should preferably not impair viral fitness and not compromise the expression of neighboring genes. Previous work by Morimoto and colleagues demonstrated that intergenic regions (IGR) between viral genes in a tail-to-tail orientation are suitable for the insertion of foreign genes into the herpes simplex virus type 1 (HSV-1) genome without compromising viral replication (27). Therefore, we decided to test the IGR between MnatCMV2 open reading frames (ORFs) M25 and m25.1. These ORFs have a tail-to-tail orientation, reducing the risk of disrupting promoter elements. The products of these two genes have not been characterized so far, but their orthologs in MCMV are known to be non-essential for viral replication in cell culture (28, 29). The MnatCMV2 BAC was modified by inserting a Kan marker into the IGR between MnatCMV2 M25 and m25.1 via *en passant* BAC mutagenesis (MnatCMV2-Kan; Fig. 6A). The Kan marker was inserted into the middle of the 95 bp IGR. Thereby, disruption of the predicted poly-A signals for both viral gene transcripts was avoided (Fig. 6A). At the same time, HA and FLAG tag sequences were added to the 3’ ends of the M25 and m25.1 ORFs, respectively, to facilitate detection of their respective protein products. In a second recombination step, the Kan marker was removed, restoring the original sequence of the M25/m25.1 IGR (reMnatCMV2-BAC). The two recombinant MnatCMV2 BACs were analyzed via restriction digest (Fig. 6B) and sequence-verified by Illumina sequencing. Infectious virus was reconstituted by transfection of MasEFs with the recombinant MnatCMV2 BACs.

**Figure 6.**
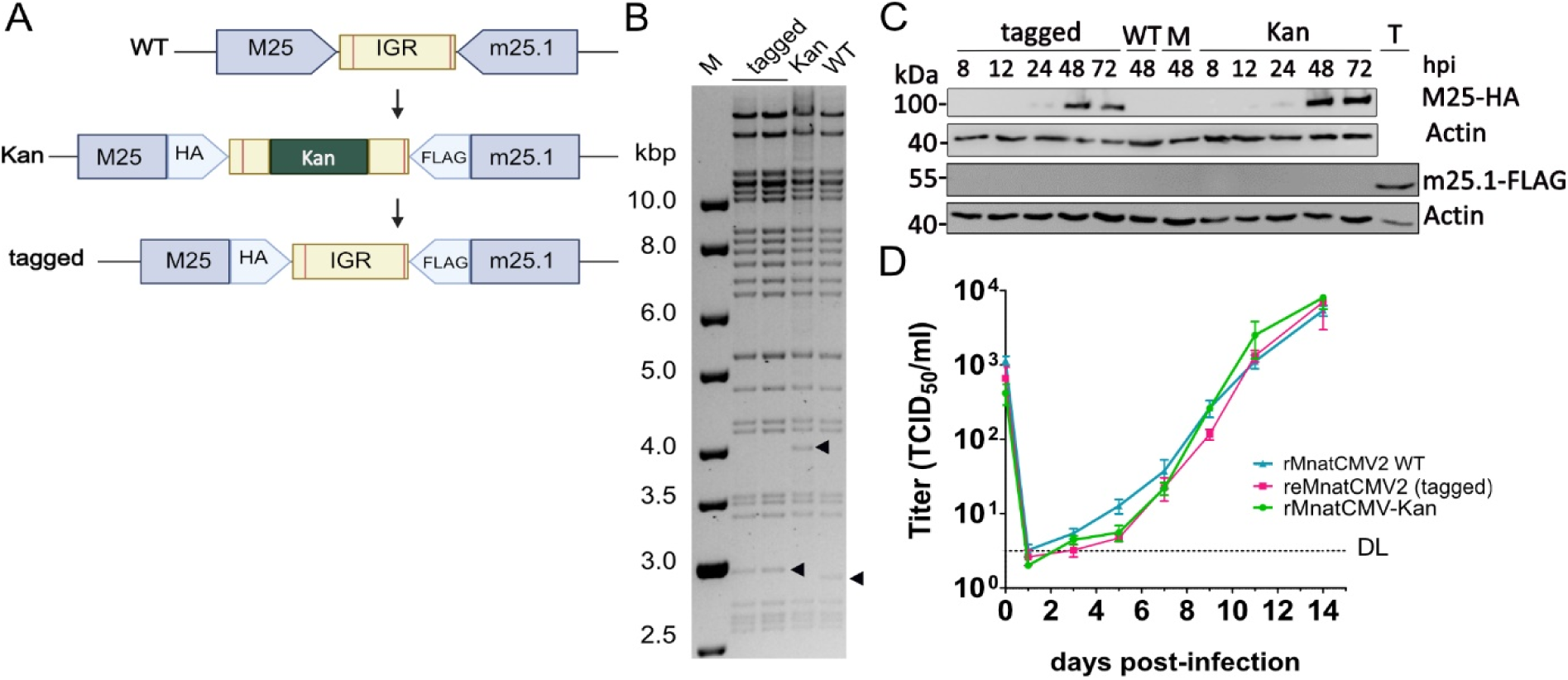
Insertional mutagenesis of the MnatCMV2 BAC. **(A)** The intergenic region (IGR) between ORFs M25 and m25.1 was modified by *en passant* mutagenesis of the MatCMV2 BAC. The M25 and m25.1 ORFs were tagged with HA and FLAG tag sequences, respectively. In a first step, a Kan cassette for selection was inserted in the middle of the IGR. In the second step, the Kan marker was removed. The vertical lines within the IGR indicate the predicted polyadenylation signals. **(B)** Restriction fragment length analysis of the mutant MnatCMV2 BACs digested with BamHI. The full-length WT MnatCMV2 BAC and the tagged mutants with or without the Kan cassette (2 clones) are shown. M, DNA size marker. Arrowheads indicate the IGR-containing band. **(C)** MasEFs were infected with the tagged viruses (MOI 0.1), and expression of the M25 and m25.1 proteins was determined by western blot analysis using anti-HA and anti-FLAG antibodies, respectively. Transfected 293T cells expressing m25.1-FLAG served as positive control (T), mock-infected (M) and parental MnatCMV2-infected cells (WT) were used as negative controls. **(D)** MasEFs were infected (MOI 0.5) with the BAC-derived rMnatCMV2 viruses. Viral titers in the supernatants (mean ±SEM of three biological replicates) were determined. DL, detection limit.

First, M25 and m25.1 protein expression was compared at different times post-infection by western blot analysis (Fig. 6C). The expression of M25-HA in infected MasEFs was comparable for both viruses with or without Kan marker, showing low protein levels at 24 hours post-infection (hpi) and increasing levels at 48 and 72 hpi. In contrast, a FLAG-tagged m25.1 protein could not be detected at the expected molecular weight at any time point, suggesting that m25.1 protein expression is very weak or absent.

Next, viral fitness was assessed by comparing multistep replication kinetics of parental and the two recombinant MnatCMV2s in MasEFs. As shown in Fig. 6D, there was no significant difference in the replication kinetics of the three viruses, indicating that insertion of a transgene between M25 and m25.1 is not detrimental to viral replication in cell culture.

## Discussion

Two recent studies have reported the identification and genetic characterization of three distinct MnatCMVs that potentially belong to genus *Muromegalovirus* in subfamily *Betaherpesvirinae* (5, 6). The present study was designed to make these viruses amenable to genetic engineering. To this end, we cloned the genomes of one isolate of each MnatCMV as a BAC using STAR in yeast. STAR cloning allows rapid and precise cloning of herpesvirus genomes as BACs in a single step. When we initially planned the cloning of the first MnatCMV genome (MnatCMV2), we were concerned that a repetitive sequence such as the TR at the genomic termini may cause unwanted genetic reassortment due to recombination of the TRs within the two homology hooks. However, through cloning of other CMV genomes performed in parallel we came to realize that short repetitive sequences, such as the 30 bp TRs of muromegaloviruses, do not compromise cloning efficiency (23). Therefore, we subsequently cloned the complete genomes of MnatCMV1 and MnatCMV3 isolates in a single step, without the need to repair an incomplete genome afterwards.

Complete sequencing of the BAC clones and the viruses reconstituted from them showed high fidelity of the technology, with no genetic differences between the parental viruses and the BAC-cloned genomes. The sequence analyses also confirmed that the YCp-BAC vector, which was placed between the genome termini, was automatically excised during virus reconstitution, probably by the viral terminase, which cuts genomes from concatemers as they are packaged into capsids (30). As the BAC-derived viruses were genetically indistinguishable from the parental viruses, it was not surprising that their replication in fibroblast and epithelial cells showed no discernible differences (Fig. 4). This achievement is important, as the preservation of replication fitness is a prerequisite for a successful transmissible vaccine.

Another important consideration was the maintenance of replication in their target host species (*M. natalensis*), but with otherwise limited host range. Although CMVs are very species-specific, there are some documented limitations and exceptions. While the RCMV Maastricht strain cannot replicate in mouse cells, MCMV can replicate in rat fibroblasts (24, 25). CMVs isolated from a field mouse and a roof rat were reported to replicate in hamster kidney cells (31, 32). All three MnatCMVs replicated to levels comparable to the parental non-recombinant viruses in two different *M. natalensis* derived cell types. However, none of the three MnatCMVs analyzed in the present study replicated to detectable titers in mouse and rat cells. A transient low-level replication of MnatCMV2 in BHK-21 cells was detected (Fig. 5). As BHK-21 cells were the only hamster cell line available, it is unclear whether MnatCMV2 is capable of replicating at very low levels in hamster cells or whether BHK-21 cells are more permissive for MnatCMV2 replication than other hamster cells. BHK-21 cells are well-known to be highly permissive for many different viruses from various host species, such as rabies virus (33), modified vaccinia virus Ankara strain (34), and herpesviruses such as HSV-1, murine gammaherpesvirus-68, and MCMV (35–37). The exceptional permissiveness of BHK-21 cells to viral infections has been explained by a lack of type I interferon signaling and an impaired activity of pattern recognition receptors (38, 39). Hence the significance of the low-level replication in BHK-21 cells remains questionable. In contrast, MnatCMV1 and MnatCMV3 did not replicate to detectable levels in BHK-21 cells (Fig. 5).

To identify a suitable transgene insertion site, we analyzed the MnatCMV2 genome for regions with non-overlapping ORFs in a tail-to-tail configuration. Such regions were previously reported to serve as ideal transgene insertion sites within the HSV-1 genome (27). We focused on the IGR between M25 and m25.1 and verified its suitability for insertion of a transgene. Insertion of a 1 kb Kan cassette did not alter viral replication kinetics. It also did not compromise expression of the M25 protein over time (Fig. 6). Even though the function of MnatCMV2 M25 is unknown, its ortholog in MCMV is known to play a role in cytoskeleton remodeling and p53 sequestration during infection (28, 40). An M25 deletion mutant replicated to lower titers *in vitro* and *in vivo* (28). By contrast, much less is known about m25.1. Its ortholog in MCMV is a member of the US22 gene family. It is nonessential for viral replication, and an m25.1-deficient virus grew to comparable titers as WT MCMV in various cell types (29). To our surprise, we were unable to detect m25.1 protein expression in MnatCMV2-infected cells (Fig. 6). This finding suggested that the m25.1 protein is expressed at extremely low levels (or not at all) in cell culture, or that the protein is highly unstable. In any case, our results indicated that the insertion of a transgene (Kan marker) into the IGR between M25 and m25.1 was neither detrimental to MnatCMV2 replication nor to the expression of the two genes (Fig. 6), regardless of their function. Based on the existing data, this site appears to be very well suited for the insertion of vaccine antigen expression cassettes within the MnatCMV genome.

In the absence of an approved LASV vaccine for humans, Lassa fever infections continue to pose a significant threat to the human population (41). Although many LASV infections only cause mild symptoms, the case-fatality rate after hospitalization still reaches 15 to 25% (42). Previous studies on *M. natalensis* populations and LASV transmission have investigated the question of which kind of intervention measures could stably reduce zoonotic transmission of LASV to humans. Field studies and mathematical models of rodent population containment and LASV transmission suggested that continuous control or rodent vaccination are the only strategies that could lead to LASV elimination (43).

In order to apply a vector-based vaccine to an adequate proportion of a hard-to-reach target population, such as *M. natalensis*, the idea of a transmissible vaccines has gained attention. Mathematical predictions for the efficiency of transmissible viral vaccines applicable to rodents predict that a CMV-based transmissible vaccine against LASV could achieve a pathogen reduction of up to 95% in just 212 days due to a relatively short infectious period and reproductive number of LASV (44, 45). As promising as such predictions may be, the development of an effective transmissible vaccine remains a great challenge. The cloning of three complete MnatCMV genomes (representing three different viral species) as BACs makes these viruses amenable to genetic modifications, which is a prerequisite for their use as vaccine vectors. Hence, this study provides a first step towards the generation of a transmissible vaccine directed against LASV in the rodent host population.

## Acknowledgments

We thank the staff of the Next Generation Sequencing technology platform at the Leibniz Institute of Virology and the MRC-University of Glasgow Centre for Virus Research Genomics and Bioinformatics Groups for technical support. Schematic figures were created using BioRender. This work was supported by the DARPA PREEMPT program (grant no. D18AC00028).

## Data availability

The complete sequences of the MnatCMV1, MnatCMV2, and MnatCMV3 BACs have been deposited in GenBank under the accession numbers PP337206, PP337207, and PP337208. All other data supporting the findings of this study are contained within the article.

## Material and Methods

### Cells and viruses

NIH-3T3 (CRL-1658), BHK-21 (CCL-10) and human embryonic kidney 293T (CRL-3216) cells were obtained from the American Type Culture Collection, and REF cells (23, 46) were provided by Sebastian Voigt (University Hospital Essen, Essen, Germany). MasKECs and MasEFs were isolated from *M. natalensis* animals bred in the animal facility of the Bernhard Nocht Institute for Tropical Medicine (BNITM) in Hamburg, Germany. MasKECs have been described previously (6, 47). MasEFs were isolated from embryos (15 -16 days of age) essentially as described previously for murine embryonic fibroblasts (48). Briefly, *M. natalensis* embryos were trypsinized for 30-60 min. The connective tissue was removed by passage through a 75 µm filter, and the cell pellet was collected and resuspended in complete Dulbecco’s modified Eagle’s medium (DMEM, Gibco) containing 10% (v/v) fetal calf serum (FCS), 100 U/ml penicillin and 100 µg/ml streptomycin (Sigma). Primary MasEFs at passage 1 were immortalized by transduction with a retroviral vector expressing the SV40 large T antigen as described (49). All cell lines were cultured at 37°C and 5% CO_2_ in DMEM, containing 5% (v/v) FCS, 100 IU/mL penicillin and 100 µg/mL streptomycin. MnatCMV1 (isolate Mnat36A E2-3), MnatCMV2 (isolate Mnat2A C3-3), and MnatCMV3 (Mnat35A A3-2) have been described previously (6). Their complete genome sequences are available in GenBank under accession numbers OP429138.1, OP429139.1 and OP429140.1.

### Plasmids

The retroviral vector for immortalization of MasKECs and MasEFs, pBABE-cLT (plasmid #1779), and pEPkan-S (plasmid #41017) were obtained from Addgene. The pcDNA3 expression plasmid and pEM7/Zeo were obtained from Invitrogen.

The YCp-BAC cloning vector, pCC1BAC-his3 (50), was kindly provided by Daniel Malouli and Klaus Früh (Oregon Health Science University, Portland, OR). It contains a bacterial F plasmid-derived replicon (BAC cassette) and a YCp replicon for propagation in bacteria and yeast, respectively. It also contains a chloramphenicol acetyltransferase and a *his3* gene for selection in *E. coli* and *S. cerevisiae*, respectively.

For STAR cloning of MnatCMV genomes, two 60 bp homology hooks were inserted into pCC1BAC-his3 between EcoRI and BamHI restriction sites as previously described (23). Briefly, two single-stranded oligonucleotides (Table 1) were combined in annealing buffer (100 mM NaCl, 10 mM Tris-HCl, pH 7.4), incubated at 95°C for 2 min, and cooled down to room temperature (RT). A complete double-stranded oligonucleotide was obtained by filling in with Klenow DNA polymerase (Thermo Fisher). After heat-inactivation, the oligonucleotide was digested with EcoRI and BamHI and inserted into the pCC1BAC-his3 vector via ligation. A ClaI restriction site was placed between the hooks for linearization of the vector and separation of the two hooks prior to cloning.

pcDNA_MnatCMV_HR, a template plasmid used for repair of the initially cloned MnatCMV2 BAC, was generated using Gibson Assembly Ultra Kit according to manufacturer’s instructions. Briefly, specific primers were produced for amplification of overlapping PCR products using MnatCMV2 vDNA and pEPkan-S as respective templates (Table 1). PCR products were purified and mixed in equal molar ratios together with linearized pcDNA3 vector (EcoRI and BamHI). After assembly, 3 µl reaction sample was used for electroporation of electrocompetent *E. coli* strain DH10B. Bacteria were plated on LB agar plates containing kanamycin and ampicillin. After incubation overnight at 37°C, DNA of bacterial colonies was analyzed via PCR.

pcDNA3 containing MnatCMV2 m25.1 ORF was cloned by ligation of linearized pcDNA3 vector (KpnI and NotI) and PCR fragment of m25.1 ORF amplified from MnatCMV BAC WT using suitable PCR primers (Table 1). After transformation of DH10B and incubation of bacteria on LB plates containing ampicillin, extracted DNA was analyzed via PCR.

### STAR cloning of MnatCMV genomes

MnatCMV DNA was isolated from virions concentrated from the supernatant of MnatCMV-infected MasEFs by proteinase K digestion, phenol/chloroform extraction, and isopropanol precipitation as described (23).

Transformation of *S. cerevisiae* strain VL6-48N (ATCC MYA-3666) and DNA isolation from transformed yeast at day 5 post-transfection was done as described previously (23). Briefly, yeast growing on yeast extract peptone dextrose (YEPD) plates were inoculated in YEPD medium until reaching an OD_600_ of 2.0 to 2.5. Cells were purified via washing and digested with zymolyase 20T (MP Biomedicals) supplemented with 2-mercaptoethanol. After 30-60 min when spheroplasting was optimal, cells were pelleted and washed in 1 M sorbitol. After transformation of spheroplast suspension with 1 µg of linearized MnatCMV DNA and 0.5 µg of linearized cloning vector, cells were plated on SD-SORB/-his agar plates at 31°C for 5 to 7 days.

Yeast colonies were screened via PCR for positive recombination. Briefly, colonies were treated with zymolyase 20T and 2-mercaptoethanol in PBS for 2 h and 30 °C. PCR was set up with 1.5 µl of template solution and appropriate primers (Table 1).

DNA from PCR-positive yeast colonies was isolated using a modified alkaline lysis protocol (23). 5-10 µL yeast-derived DNA solution was used to transform *E. coli* strain DH10B by electroporation using a BioRad Gene Pulser II. Bacteria were plated on LB agar plates containing chloramphenicol and incubated overnight at 37°C.

### BAC DNA purification and analysis

BAC DNA was isolated from 500 mL LB culture using a NucleoBond Xtra Midi kit (Macherey-Nagel) according to the manufacturer’s protocol for low-copy plasmids. For restriction fragment analysis, 1 µg BAC DNA was digested with restriction enzymes and separated by electrophoresis on a 0.6% agarose/0.5×TBE (45 mM Tris, 45 mM boric acid, 1 mM Na_2_EDTA, pH 8.0) gel. BAC-cloned viral genomes were sequence-verified by Illumina sequencing and compared to the parental viral genome sequence. Approximately 100 ng DNA was sheared in an S220 focused-ultrasonicator (Covaris) to fragments of approximately 450 bp. Sequencing libraries were produced by conducting seven PCR cycles using a KAPA LTP library preparation kit (Roche Sequencing and Life Science) with NEBNext multiplex oligos for Illumina (New England Biolabs). The sequences were *de novo* assembled using SPAdes genome assembler v3.15.5 and aligned to the respective reference genomes using Bowtie2 as previously described (23). The complete MnatCMV BAC sequences have been deposited in GenBank (accession numbers PP337206, PP337207, and PP337208).

### MnatCMV stocks and replication kinetics

To reconstitute MnatCMV from cloned BAC DNA, MasEFs were transfected with purified BAC DNA using GenJet (SignaGen) transfection reagent as described previously (23). To compare the replication kinetics of parental MnatCMV and reconstituted BAC clones, virus stocks were prepared and titrated essentially as described (23). Briefly, MasEFs were infected at a MOI of 0.1. After 10 d, 200 ml virus-containing supernatant was centrifuged at 27,000 × *g* for 3 h. At the same time, infected cells were harvested by scraping and homogenized using a Dounce homogenizer. Cell homogenate and viral pellet were mixed and resuspended in 3-6 ml of DMEM, aliquoted, and stored at -80°C. Titration of virus stocks was performed using the 50% tissue culture infectious dose (TCID_50_) method (51) and calculated using the Spearman-Kärber formula.

For the analysis of MnatCMV replication kinetics on MasEFs, MasKECs, BHK-21 cells, REFs, or NIH-3T3 cells were seeded 1 d prior to infection in 6-well plates (2×10^4^ cells per well). Four hours after infection of the cells at the indicated MOI, the inoculum was removed, the cells were washed twice with PBS, and fresh medium was added. Supernatants were collected every 2 d for 2 weeks and stored at -80°C. Virus titers were determined by titration of the supernatants using the TCID_50_ method.

### Western Blot

To analyze viral protein expression, cells grown in 48-well plates were infected at an MOI of 0.1. At various times after infection, cells were lysed in SDS-PAGE sample buffer (125 mM Tris pH 6.8, 4% SDS, 20% glycerol, 10% 2-mercaptoethanol, 0.002% bromophenol blue). For m25.1-FLAG control, 293T cells were transfected with pcDNA3 vector containing the m25.1 ORF using GeneJet transfection reagent, and cell lysates were generated 24 h post transfection using SDS-PAGE sample buffer. Insoluble material was removed by centrifugation. Equal volumes of sample were subjected to SDS-PAGE followed by transfer to a nitrocellulose membrane (Amersham). Proteins of interest were detected with protein-specific primary antibodies and HRP-coupled secondary antibodies by enhanced chemiluminescence (Amersham) supplemented with 10% of Lumigen TMA-6 (Bioquote). Monoclonal antibodies against the following proteins and epitopes were used: β-actin (AC-74; Sigma); HA (3F10; Roche); FLAG (F7425; Sigma). All western blots shown are representative of three or more experiments.

### *En passant* mutagenesis

Introduction of genetic alterations into the BAC-cloned MnatCMV2 genome, as well as repair of missing genome ends in the initially STAR cloned-MnatCMV2 BAC, was done by *en passant* mutagenesis (52). Briefly, MnatCMV BACs were transferred to *E.coli* strain GS1783, which contains a temperature-inducible Red recombinase system as well as an L-arabinose inducible I-SceI restriction enzyme. Recombination-proficient electrocompetent bacteria were prepared as described (52) Repair of the initially incomplete MnatCMV2 BAC was done in two separate steps (i.e., repair of LJ and RJ). For the first step, a PCR product containing the missing part of LJ, two TRs and an I-Sce linked Kan resistance cassette flanked by 80 bp of duplications was used. pcDNA_MnatCMV_HR plasmid served as the template for amplification. Primers additionally contained homologies for H1 and pCC1BAC-his3 present in the MnatCMV2 BAC to facilitate homologous recombination. After transformation with the purified PCR product, bacteria were positively selected using chloramphenicol and kanamycin. The second recombination step released the Kan cassette via the 80 bp duplications. BAC DNA of kanamycin sensitive clones was verified using restriction digestion. The RJ region adjacent to H2 was repaired accordingly using respective primers (Table 1).

MnatCMV2 BAC mutagenesis of the IGR between the M25 and m25.1 ORFs was performed as follows. First, the complete IGR was replaced by a zeocin resistance cassette and the M25/m25.1 ORFs were modified to contain C-terminally located HA- and FLAG-tags, respectively. Mutagenesis was done using one primer pair with homologies to M25/m25.1 ORFs. pEM7/Zeo served as template plasmid. After transformation, bacteria were positively selected using chloramphenicol and zeocin. Correct recombination in the resulting MnatCMV2-BAC-Zeo was verified via PCR. Next, the IGR sequence was ordered as a gBlock from Integrated DNA Technologies (IDT) together with homologies to M25-HA and m25.1-FLAG ORFs (Table 1) and was cloned into pcDNA3 via ligation. An I-Sce linked Kan resistance cassette flanked by 40 bp of duplication homologous to the IGR was inserted into the IGR cloned into pcDNA3. The M25-HA-IGR-Kan-IGR-FLAG-m25.1 construct was excised from pcDNA3 via restriction digestion and used for homologous recombination with MnatCMV2-BAC-Zeo. This resulted in replacement of the Zeo marker by the IGR carrying the I-Sce/Kan resistance cassette. A second recombination step released the Kan marker via the 40 bp duplication within the IGR. The structures of both Kan-resistant and Kan-sensitive BAC clones (MnatCMV-Kan and reMnatCMV-BAC, respectively) were verified by restriction digestion.

### Ethical Statement

MasKECs and MasEFs were isolated at LIV from *M. natalensis* rodents according to the recommendations and guidelines of the Federation for Laboratory Animal Science Associations (https://felasa.eu/) and the Gesellschaft für Versuchstierkunde (Society of Laboratory Animal Science; https://www.gv-solas.de/). The procedures were approved by the institutional review board and local authorities (Behörde für Gesundheit und Verbraucherschutz, Amt für Verbraucherschutz, Freie und Hansestadt Hamburg, reference 2018-08-16-02).

